# Integrated spatial models foster complementarity between monitoring programs in producing large-scale bottlenose dolphin indicators

**DOI:** 10.1101/2021.02.01.429097

**Authors:** Valentin Lauret, Hélène Labach, Daniel Turek, Sophie Laran, Olivier Gimenez

## Abstract

Over the last decades, large-scale ecological projects have emerged that require collecting ecological data over broad spatial and temporal coverage. Yet, obtaining relevant information about large-scale population dynamics from a single monitoring program is challenging, and often several sources of data, possibly heterogeneous, need to be integrated. In this context, integrated models combine multiple data types into a single analysis to quantify population dynamics of a targeted population. When working at large geographical scales, integrated spatial models have the potential to produce spatialised ecological estimates that would be difficult to obtain if data were analysed separately.

In this paper, we illustrate how spatial integrated modelling offers a relevant framework for conducting ecological inference at large scales. Focusing on the Mediterranean bottlenose dolphins (*Tursiops truncatus*), we combined 21,464 km of photo-identification boat surveys collecting spatial capture-recapture data with 24,624 km of aerial line-transect following a distance-sampling protocol. We analysed spatial capture-recapture data together with distance-sampling data to estimate abundance and density of bottlenose dolphins. We compared the performances of the distance sampling model and the spatial capture-recapture model fitted independently, to our integrated spatial model.

The outputs of our spatial integrated models inform bottlenose dolphin ecological status in the French Mediterranean Sea and provide ecological indicators that are required for regional scale ecological assessments like the EU Marine Strategy Framework Directive. We argue that integrated spatial models are widely applicable and relevant to conservation research and biodiversity assessment at large spatial scales.

## Introduction

Macro-institutions get increasingly involved in large-scale programs for biodiversity conservation over regional and continental areas. Whether these policies aim at assisting governments (e.g., the Intergovernmental Science-Policy Platform on Biodiversity and Ecosystem Services), or at implementing environmental management such as the European Union directives (Habitat Directive, 92/43/EEC, or Marine Strategy Framework Directive, MSFD, 2008/56/EC), conducting large-scale ecological monitoring is required to establish conservation status of targeted species and ecosystems, and to inform decision-making.

For biodiversity management decisions, conservation sciences require assessing the ecological status of species and ecosystems, which democratized the call for ecological indicators (Buckland, Magurran, et al., 2005; Nichols & Williams, 2006). An ecological indicator can be defined as a metric reflecting one or more components of the state of ecological systems. An ecological indicator can either be measured directly or result from the simplification of several field-estimated values (Niemi & McDonald, 2004). The Marine Strategy Framework Directive referred to abundance/density of targeted species (e.g. seabirds, cetaceans) as ecological indicators to fulfil for national reporting. At large spatial scales, logistical and financial constraints often prevent a detailed coverage of the targeted population using a single collection effort, and different monitoring programs coexist (Lindenmayer & Likens, 2010; Zipkin & Saunders, 2018; Isaac et al., 2019). The multiplication of monitoring programs over the same conservation context has fostered the development of statistical models that can estimate ecological quantities while accommodating several, possibly heterogeneous, datasets (Besbeas et al., 2002; Miller et al., 2019; Isaac et al., 2019; Zipkin, Inouye, & Beissinger, 2019; Farr et al., 2020). Integrating data from several monitoring protocols can give complementary insights on population structure and dynamics (Schaub & Abadi, 2011), increase space and time coverage of the population (Schaub & Abadi, 2011; Zipkin, Inouye, & Beissinger, 2019), and produce more precise ecological estimates (Isaac et al. 2019; Lauret et al. 2021; Farr et al. 2020).

A recurrent objective of ecological monitoring programs is to estimate population abundance and density (Williams, Nichols, & Conroy, 2002), for which distance sampling (DS, Buckland et al., 2005) and capture-recapture (CR, Williams et al. 2002) methods are widely used. Abundance reflects the estimated number of animals in a specified area while density is a spatialised estimate that reflects the number of animals per unit area. DS and CR methods have strengths and weaknesses in relation to logistical and practical issues (Hammond et al., 2021). DS methods can cover large areas at a reasonable cost (e.g. line transect monitoring), while CR monitoring programs can be costly to develop at large spatial scales because more sampling effort is required over a longer time period to recapture individuals (Hammond et al., 2021). Even when estimating abundance over the same study area, DS and CR do not estimate exactly the same quantity (Calambokidis & Barlow, 2004; Crum, Neyman, & Gowan, 2021). DS methods estimate abundance within a study area at the time of the survey. CR methods are based on individuals sampling and estimate the number of animals that were present in the study area during the time of the monitoring (Calambokidis & Barlow, 2004). CR methods encapsulate longer temporal extent because multiple sampling occasions are needed to build CR histories (Williams, Nichols, & Conroy, 2002). However, when data are collected over the same monitoring period and if animals do not move in and out of the study area during that period, CR and DS provide consistent estimates. Recent modelling tools have emerged to integrate both DS and CR methods into integrated population models (Kéry & Royle 2020). DS and spatial CR methods (SCR) allows accounting for spatial variation in abundance and density (Camp et al., 2020; Miller et al., 2013; Royle et al., 2014), possibly at large scales (Bischof et al., 2020). The extension to integrated spatial models has been proposed to account for spatial variation in abundance and demographic parameters while analysing jointly DS data and SCR data (Chandler et al., 2018). Integrated modelling holds promise for species occurring over large areas that are likely to be the target of multiple monitoring protocols. Besides, working at large geographical scales requires encapsulating spatial dimensions in the estimation of ecological quantities. Integrated spatial models allow to assess spatialised ecological inference, e.g density of individuals. To date, integrated spatial models have been developed and used on open populations to estimate temporal variation in population dynamics and vital rates such as survival and recruitment (Chandler & Clark, 2014; Chandler et al., 2018; Sun, Royle, & Fuller, 2019). These applications rely on long-term datasets that are not always compatible with conservation objectives. In many cases, ecological information is needed quickly, and data to investigate temporal variation are unavailable (Nichols & Williams, 2006; Lindenmayer & Likens, 2010). Consequently, ecological inference is often restricted to closed-population indicators (e.g. abundance or population size, density or spatial repartition of the population, distribution or spatial extent of a population). When the temporal resolution of monitoring programs does not allow to quantify population dynamics, we argue that an application of integrated spatial models to closed populations can be useful in numerous ecological contexts to deal jointly with existing monitoring programs and assess abundance and density.

In this paper, we build an integrated spatial model and demonstrate the relevance of combining DS and SCR to build large-scale ecological indicators. We consider the monitoring of common bottlenose dolphins (*Tursiops truncatus*) that are considered as “vulnerable” by the IUCN Red List in the North-Western Mediterranean Sea (IUCN, 2009). The protected status of bottlenose dolphins within the French seas (listed on Annex II of the European Habitats Directive) led to the development of specific programs to monitor Mediterranean bottlenose dolphins within the implementation of the European Marine Strategy Framework Directive, which requires assessing the conservation status of this species every 6 years over the large extent of the French Mediterranean Sea (Authier et al. 2017). Increasing efforts are dedicated to develop monitoring programs in the Marine Protected Areas (MPA) network that mainly implement photo-identification protocols locally, while governmental agencies perform large-scale line-transect programs to monitor marine megafauna and fisheries. Hence, multiple data sources coexist about bottlenose dolphins in the French Mediterranean Sea. In this paper, we analysed DS data collected by aerial line-transect surveys over a large area covering coastal and pelagic seas (Laran et al., 2017), which we combined with SCR data collected by a photo-identification monitoring program restricted to coastal waters (Labach et al., 2021). We compared the abundance and density of bottlenose dolphins estimated from DS model, SCR model, and integrated spatial models to highlight the benefits of the integrated approach in an applied ecological situation. We discussed the promising opportunities of using integrated spatial models in the context of marine monitoring planning in the French Mediterranean. Eventually, we underlined the conservation implications of using such a model at a wider extent to make the best use of available datasets.

## Methods

### Monitoring bottlenose dolphins in the French Mediterranean Sea

Common bottlenose dolphins (*Tursiops truncatus*) occur over large areas throughout the Mediterranean Sea. Because monitoring elusive species in the marine realm is complex, multiple monitoring initiative have emerged to collect data about bottlenose dolphins in the French Mediterranean Sea. In the context of the Marine Strategy Framework Directive, the French government implemented large-scale aerial transects to monitor marine megafauna (Laran et al., 2017). However, the large spatial coverage of the aerial monitoring is impaired by the low resolution of such data (i.e. 1 campaign every 6 years). Then, to collect detailed data, the French agency for biodiversity funded a photo-identification monitoring program to investigate the ecological status of the bottlenose dolphins in the French Mediterranean Sea. This coastal boat photo-identification monitoring has been performed between 2013 and 2015. Coastal photo-identification monitoring represents a promising opportunity to produce high resolution information because data can be collected routinely by Marine Protected Areas at high time frequency.

### Study area and datasets

We focused on an area of 255,000 km^2^ covering the North-Western Mediterranean Sea within which we considered two monitoring programs about bottlenose dolphins. We used SCR data from at-sea boat surveys over 21,464 km of the French continental shelf. Observers performed monitoring aboard small boats to locate and photo-identify bottlenose dolphins all year long between 2013 and 2015, always at constant speed and with three observers. Taking pictures of the dorsal fin of each individual in the group makes possible the construction of detection history and hence the analysis of the population through capture-recapture methods (Labach et al., 2021). Boat surveys were restricted to the coastal waters of France and adopted a search-encounter design covering approximatively all the continental shelf every 3 months. We divided the duration of the monitoring programs into 8 equal sampling occasions that length for 3 months each, following previous analysis by Labach et al., (2021). We also used DS data that were collected during winter and summer aerial line-transect surveys covering 24,624 km of both coastal and pelagic NW Mediterranean Sea between November 2011 to February 2012 and May to August 2012 (Laran et al., 2017). Two trained observers collected cetacean data following a DS protocol (i.e. recording species identification, group size, declination angle). Aerial surveys were conditional on a good weather forecast.

We divided the study area in 4356 contiguous grid-cells creating a 5’x5’ Mardsen grid (WGS 84). To model density of individuals, we used depth as an environmental covariate, which is expected to have a positive effect on bottlenose dolphins’ occurrence (Bearzi, Fortuna, & Reeves, 2009; Labach et al., 2021). To estimate the sampling effort of aerial and boat surveys, we calculated the transect length (in km) prospected by each monitoring protocol within each grid-cell during a time period. Typically, entire transects are split into segments as they overlap multiple grid-cells (Miller et al., 2013). Sampling effort was therefore cell and occasion-specific in the case of the SCR model, and cell specific for the DS model. Sampling effort ranged from 0.047 km to 308 km per grid-cell and per occasion for the photo-id dataset, and from 1.33 to 54560 km per grid-cell for the aerial line transect dataset. We used subjective weather condition recorded by plane observers during the line transect protocols as a discrete variable ranging from 1 to 8. Good weather condition was expected to be positively related to the detection probability.

### Spatial integrated models for closed populations

To integrate DS and SCR data, we used the hierarchical model proposed by Chandler et al. (2018). However, while initially developed for open populations and due to the lack of temporal depth in our datasets, we adapted the model to estimate abundance and density without accounting for demographic parameters (Fig 1). We performed closed population estimation of bottlenose dolphin density over the 2011-2015 period, assuming that (1) the population was demographically closed during the study period, (2) all individuals were correctly identified at each capture occasion and marks were permanent during the sampling period, (3) no migratory events occurred during the sampling period. Although being strong assumptions, bottlenose dolphin deaths and recruitments between 2011 and 2015 were likely small considering the long life cycle of bottlenose dolphins (Bearzi, Fortuna, & Reeves, 2009; Hammond et al., 2021). Besides, Western Mediterranean bottlenose dolphin population is clustered into coastal subunits, hence we neglected migration events and movements that can occur between “French” resident groups and other populations, e.g. offshore, Spanish, Italian or Atlantic groups (Louis et al., 2014; Carnabuci et al., 2016).

**Figure 1:**
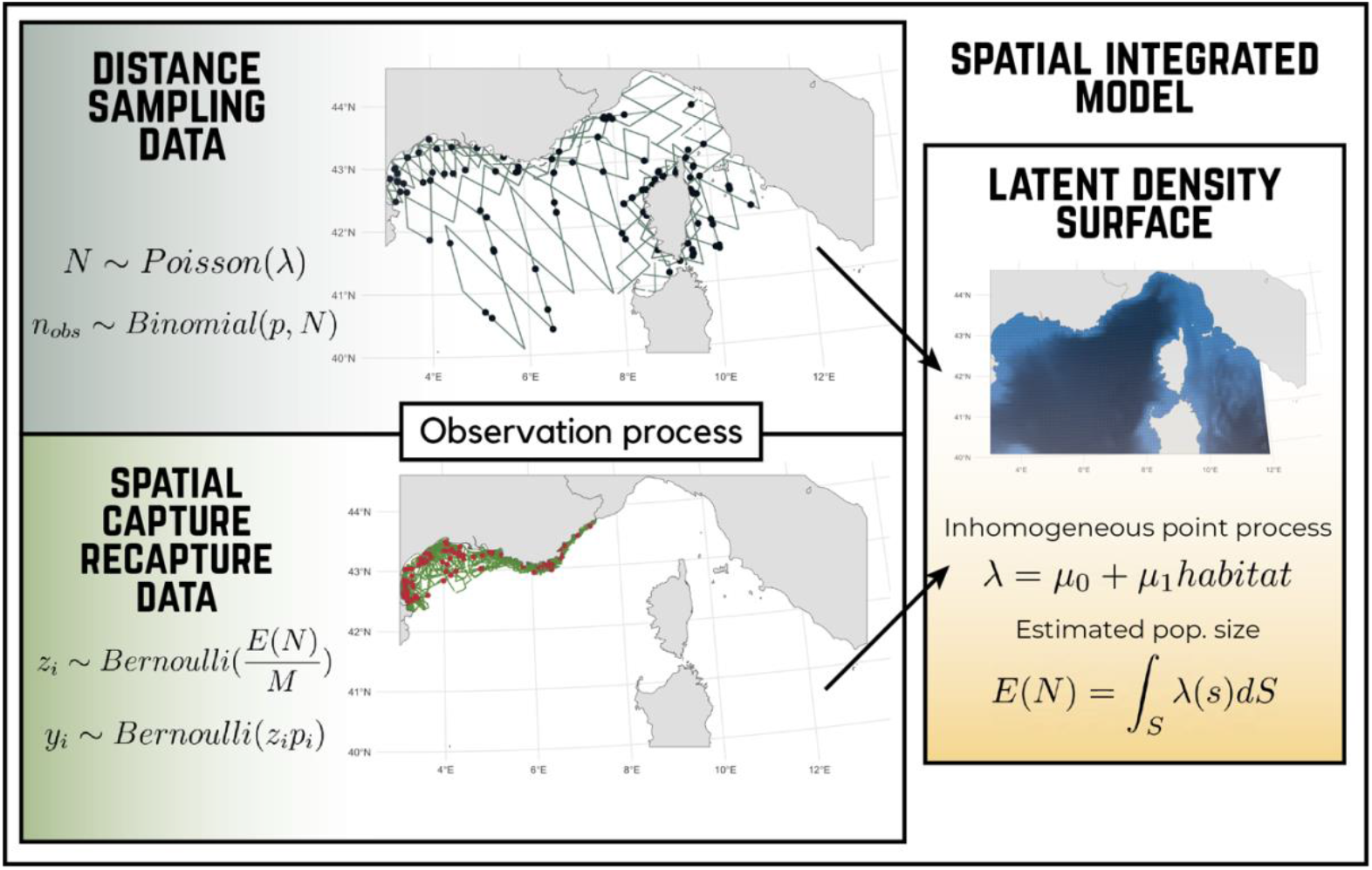
Graphical description of the Spatial Integrated Model (SIM) that combines Spatial Capture Recapture (SCR), and Distance Sampling (DS). The SIM is a hierarchical model with three processes: i) latent population size E(N) and density λ informed by an inhomogeneous point process, ii) DS observation process that link the line-transect dataset to the latent density surface, iii) SCR observation process that links the detection histories to the latent density. The observation process is stochastic according to detection probability. For DS model, the observed group size n_obs_ is a Binomial draw in the latent abundance N at the sampled grid-cell. For SCR model, observing an individual *i* is a Bernoulli draw with a detection probability *p*_*i*_. Through the data augmentation process with a hypothetical population size M, the probability an individual *i* belongs to the study population is the result of a Bernoulli draw of probability E(N)/M.

We structure our integrated spatial model around two layers with i) an ecological model that describes the density of individuals based on an inhomogeneous point process (*Spatial abundance* section below), and ii) two observation models that describe how the DS and SCR data arise from the latent ecological model (*Capture-recapture data* and *Distance-sampling data* sections below).

### Spatial abundance

For the ecological model, we use a latent spatial point process modelling the density of individuals and the overall abundance. Over the study area *S*, an intensity function returns the expected number of individuals at location *s* in *S*. Here, *s*, represents an arbitrary point in the study area *S*. To account for spatial variation, we model the latent density surface as an inhomogeneous point process. For every location *s* in the study area *S*, the expected abundance λ is written as a log-linear function of an environmental covariate, say habitat:

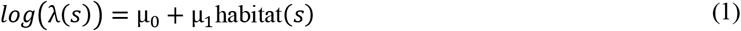

where parameters to be estimated are *µ*_*0*_ and *µ*_*1*_ respectively the density intercept and the regression coefficient of the environmental covariate. For simplicity, we use depth as a habitat covariate possibly influencing bottlenose dolphin density, and explore a linear relationship between density and depth. The effect of habitat covariates could be further explored (e.g. by considering other covariates such as sea surface temperature or prey availability, or by accounting for non-linear effects). Then, the estimated population size is derived by integrating the intensity function over the study area:

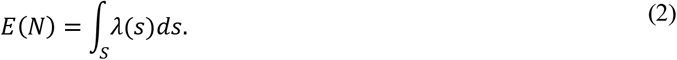

As we discretized the study area, we estimated *λ*_*j*_ the intensity process describing density for each grid-cell *j* with *j* = 1, …, *J* = 4356, hence 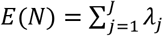. The latent ecological process defined by Eq. 1 is an inhomogeneous point process that is common to both the SCR and DS models. SCR and DS data are linked to density λ and informed the parameters of Eq. 1. To account for unseen individuals, we used the data augmentation technique and augmented the observed datasets to reach M = 20,000 individuals (Royle & Dorazio, 2012). Each individual *i* is considered being (*z*_*i*_ *=* 1) or not (*z*_*i*_ *=* 0) a member of the population according to a draw in a Bernoulli distribution of probability Ψ, with *z*_*i*_∼Bernoulli(*ψ*) where Ψ is the probability for individual *i* to be a member of the population, with *ψ* = *E*(*N*)/*M* and 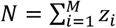.

### Capture-recapture data

To link capture-recapture data with the ecological process, we built a SCR model (Royle et al., 2014). Detection history of individuals were collected over T = 8 sampling occasions and capture locations were recorded. Grid-cells *j* in which sampling effort was positive during an occasion were considered as active detectors for this sampling occasion, hence reflecting that animals could be observed. We stored observations in a three-dimensional array *y* with *y*_*ijt*_ indicating whether individual *i* was captured at grid-cell *j* during sampling occasion *t*. We assume that observation *y*_*ijt*_ is an outcome from a Bernoulli distribution with capture probability *p*_*ijt*_, *y*_*ijt*_∼Bernoulli(*p*_*ijt*_ *z*_*i*_). We model capture probability with a half-normal detection function 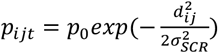 where *d*_***ij***_ is the Euclidian distance between the activity center of individual *i* and the grid-cell *j, σ*_*SCR*_ is the scale parameter of the half-normal function, and *p*_*0*_ is the baseline encounter rate (Royle et al., 2014). We accounted for spatial and temporal variation in the detection probability through the baseline detection rate p_0_ that we modelled as a logit-linear function: logit(*p*0_*jt*_) = *δ*_0_ + *δ*_1_*E*_*jt*_. When the sampling effort E_jt_ is null, we fixed *p0*_*ijt*_ to 0.

The locations of activity center inform the density of individuals λ. For each individual *i* belonging to the sampled population, its activity center is assigned as the result of a multinomial draw in the predicted density in each grid-cell of the study area.

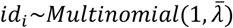

where *id*_*i*_ is the activity center of individual *i*, and 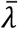 represent the vector of the predicted density in each cell of the study area. Due to the computational burden to sample the 4356 grid-cells, we mimicked the multinomial distribution through the “zeros trick” (see R codes for details). We considered that activity centers did not change between sampling occasions.

### Distance-sampling data

To accommodate distance data, we built a hierarchical DS model (Kéry & Royle, 2016). We model the DS data conditional on the underlying density surface defined by Eqs (1) and (2). We considered two sampling occasions *t*_*ds*_ as some transects were replicated. We assume that the probability of detecting dolphins is a decreasing function of the perpendicular distance between the transect and dolphin group. Because distance may not be estimated with perfection by observers, we discretized the distance of observation in B distance bins. Then, 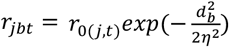, where η is the scale parameter of the half-normal function, and *r*_*0(j)*_ is the probability of detection in the grid-cell *j*, and *d*_*b*_ is the observation distance between the flight transect and bin *b* where the detection occurred. The distance class *d*_*j*_ of the observed data at grid-cell *j* is modelled as a multinomial/categorical draw

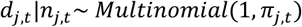

with *π*_*j,t*_ the vector of length B storing the detection probabilities in each bin *b* at grid-cell *j*. The *b*^th^ index being *π*_*j,b,t*_ = *r*_*j,b,t*_/(∑_*b*_ *r*_*j,b,t*_).

We account for spatial variation in the baseline detection rate of the detection function modelling *r*_0(*j,t*)_ as a log-linear function of weather condition *W*_*j,t*_ in grid-cell *j* during sampling occasion *t*:

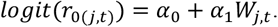

Besides, aerial surveys only sampled a fraction of the total area of each grid-cell (Appendix 1). We calculated *S*_*j,t*_ the proportion of the grid-cell effectively sampled by aerial surveys considering a 1200 m wide annulus around the transect. We assumed that density within each grid-cell was uniform and remained constant across the sampling period. Then, *N*_*j,t*_ the number of individuals sampled by aerial surveys in each grid-cell *j* during sampling occasion *t* is Poisson distributed with *λ*_*j*_ being the density of individuals predicted by the point process in grid-cell *j* restricted to the proportion of grid-cell sampled, *S*_*j,t*_.

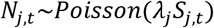

Then, *n*_*j,t*_ the observed group size detected at grid-cell *j* during sampling occasion *t*, is given by a Binomial draw in the expected number of sampled individuals, *N*_*j,t*_ with probability the sum of *r*_*j,b,t*_, the detection probabilities within each bin *b* of grid-cell *j* during sampling occasion *t*.

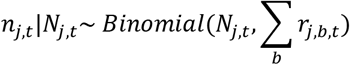

### Bayesian implementation

To highlight the benefit of integrating data for the estimation of bottlenose dolphin density, we compared i) the output of the spatial DS model, ii) the SCR model, and iii) the integrated spatial model.

We ran all models with three Markov Chain Monte Carlo chains with 100,000 iterations each in the NIMBLE R package (de Valpine et al., 2017). We checked for convergence calculating the *R-hat* parameter (Gelman et al., 2013) and reported posterior mean and 80% credible intervals (CI) for each parameter. We considered as important the effect of a regression parameter whenever the 80% CI of its posterior distribution did not include 0. We also calculated the predicted density of bottlenose dolphins (i.e. *λ*). Data and codes are available on GitHub (*https://github.com/valentinlauret/SpatialIntegratedModelTursiops*).

## RESULTS

We detected 536 dolphins through aerial surveys clustered in 129 groups. We identified 927 dolphins over 1707 detections in photo-identification surveys, out of which 638 dolphins were captured only once (68%), 144 were captured twice (15.5%), 149 were captured 3 times and up to 8 times for one individual. The maximum distance between two sightings of the same individual was 302 km, with one individual detected twice during the same sampling occasion at 115 km distance.

We estimated 2451 dolphins (2337; 2566) with integrated spatial model over the study area (Table 1), 11531 dolphins (10132; 12997) with the DS model and 1834 dolphins (1745; 1926) with the SCR model (Table 1). Density intercepts of integrated spatial model (µ_0_= -0.85 (−0.90; -0.79)) and SCR model (µ_0_= -1.18 (−1.81; -1.07)) were lower than intercept of DS model (µ_0_= 0.95 (0.82; 1.08)).

**Table 1:**
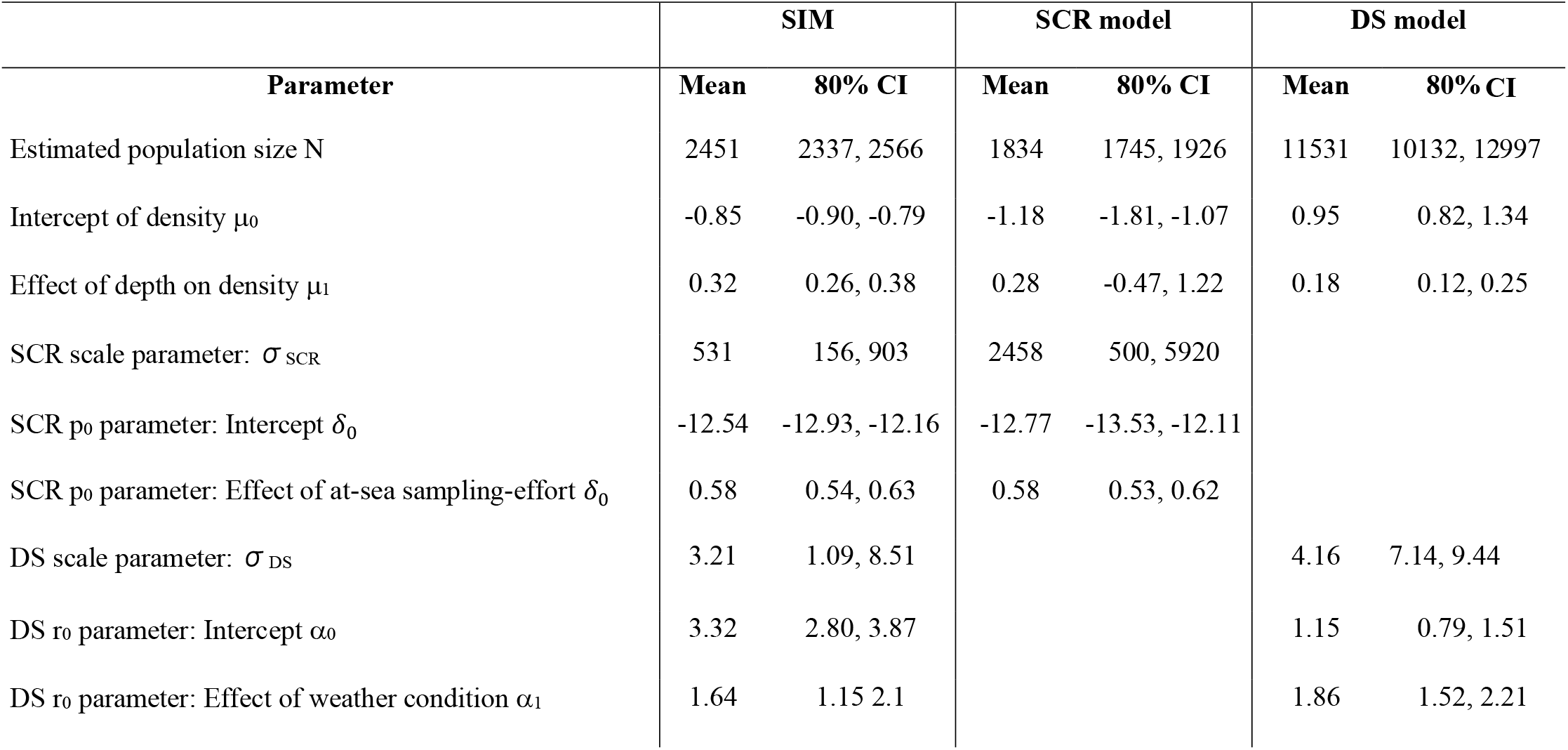
Parameter estimates for the spatial integrated model (SIM), spatial capture-recapture (SCR) model, and distance-sampling (DS) model. For each parameter, we display the posterior mean and its 80% credible interval (CI).

DS model estimated a positive effect of shallow waters (µ_1_= 0.18 (0.12; 0.25), Table 1) similar to the effect estimated by the integrated spatial model (µ_1_= 0.32 (0.26; 0.38), Table 1). However, the SCR model did not detect an effect of depth on density (µ_1_= 0.28 (−0.47; 1.22), Table 1). Then, both integrated and DS models predicted higher densities of bottlenose dolphins in the coastal seas than in the pelagic seas, whereas the SCR model predicted no effect of depth on dolphin density.

Boat sampling effort exhibited a positive effect on detection probability for both the SCR model (β_1_= 0.58 (0.53; 0.62)) and the integrated spatial model (β_1_= 0.58 (0.54; 0.62), table 1). For the integrated spatial model and the DS model, the detection probability increased when the weather condition improved (integrated spatial model: α_1_= 1.64 (1.15; 2.10), DS: α_1_= 1.86 (1.52; 2.21), Table 1).

## DISCUSSION

### Integrated spatial model benefits from both distance sampling and capture-recapture data

With our integrated spatial model, we estimated bottlenose dolphin abundance within the range of what was found in previous studies in nearby areas (Gnone et al., 2011; Lauriano et al., 2014), and found that densities were more likely to be higher in coastal areas (Bearzi, Fortuna, & Reeves, 2009). A striking result was the higher abundance estimated by DS compared to abundance estimated by the integrated and SCR models, which estimates were also found in previous studies analysing the same datasets in isolation. Using CR data only, Labach et al. (2021) estimated 2350 dolphins (95% confidence interval: 1827; 3135) inhabiting the French continental coast where our integrated model predicted 2451 dolphins (95% confidence interval: 2306; 2602). Analysing DS data, Laran et al., (2017) estimated 2946 individuals (95% confidence interval: 796; 11,462) during summer, and 10,233 (95% confidence interval: 4217; 24,861) during winter where our DS model estimated 11,531 (95% confidence interval: 9784; 13,478) all year long. Recent aerial campaigns performed in 2018-2019 on the same study area and following the same distance sampling protocol do not suggest seasonal difference in bottlenose dolphins abundance (Laran et al., 2021).

We see several reasons that might explain the discrepancy in estimates obtained from SCR and DS models. First, although the Mediterranean bottlenose dolphins population is clustered in coastal sub-units (Carnabuci et al., 2016), groups can be encountered offshore (Bearzi, Fortuna, & Reeves, 2009). In the DS dataset, large dolphin groups were detected in the pelagic seas at the extreme south of sampling design (Appendix 1). These groups could either be i) occasional pelagic individuals belonging to coastal populations and that are mainly resident outside our study area (e.g. Balearic, South-Western Sardinia), or ii) resident pelagic populations that are not sampled by coastal photo-id surveys (Louis et al., 2014). Second, SCR data were restricted to the French continental coast and did not sample dolphin populations that exist elsewhere in the study area, e.g. in Corsica, Liguria, and Tuscany (Carnabuci et al., 2016). Despite this geographic sampling bias in the capture-recapture data, SCR models should predict the existence of Corsican and Italian populations if the relationship between density and habitat in Eq (1) was correct and consistent throughout the study area. Predicting abundance outside the range of the data used could lead to biased estimates if the habitat-density relation is not correctly specified (A. Lee-Yaw et al., 2021; Hammond et al., 2021). As the photo-id surveys did not sample greater depths, our SCR model is likely to underestimate abundance because the relation linking dolphin density to depth was not correctly specified. Thus, we emphasized the relevance of aerial surveys that collected data in the pelagic seas, which helps to quantify the habitat-density relationship. To perform detailed analysis of the NW Mediterranean bottlenose dolphin populations, one should consider additional environmental covariates to better capture spatial variation in density (e.g., sea surface temperature, distance to coast, or 200m contour, Lambert et al. 2017). Besides, because Sardinian and Balearic populations, and offshore groups can be sampled in the aerial surveys, the DS model drives upward abundance compared to the SCR model that is unlikely to account for animals that are members of the Southern neither the Eastern or offshore populations.

Overall, both DS and SCR data affected the estimates of the integrated spatial model. Using SCR data brought more information about population size (e.g. more detections, more individuals) than the DS data to inform the intercept of density (µ_0_), making the integrated spatial model abundance estimate closer to the SCR model estimate (Table 1, Fig. 2). However, the DS data that were collected throughout the range of the habitat predictor informed the slope of the inhomogeneous point process (µ_1_), i.e. the effect of depth on dolphin density. Then, in the integrated spatial model, the SCR data informed the estimated population size and the DS data informed spatial repartition of individuals by correcting for the geographic sampling bias in the SCR data. The integrating approach helped to reduce the sampling limitations of each dataset and can improve the ecological inference as illustrated here about bottlenose dolphins.

**Figure 2:**
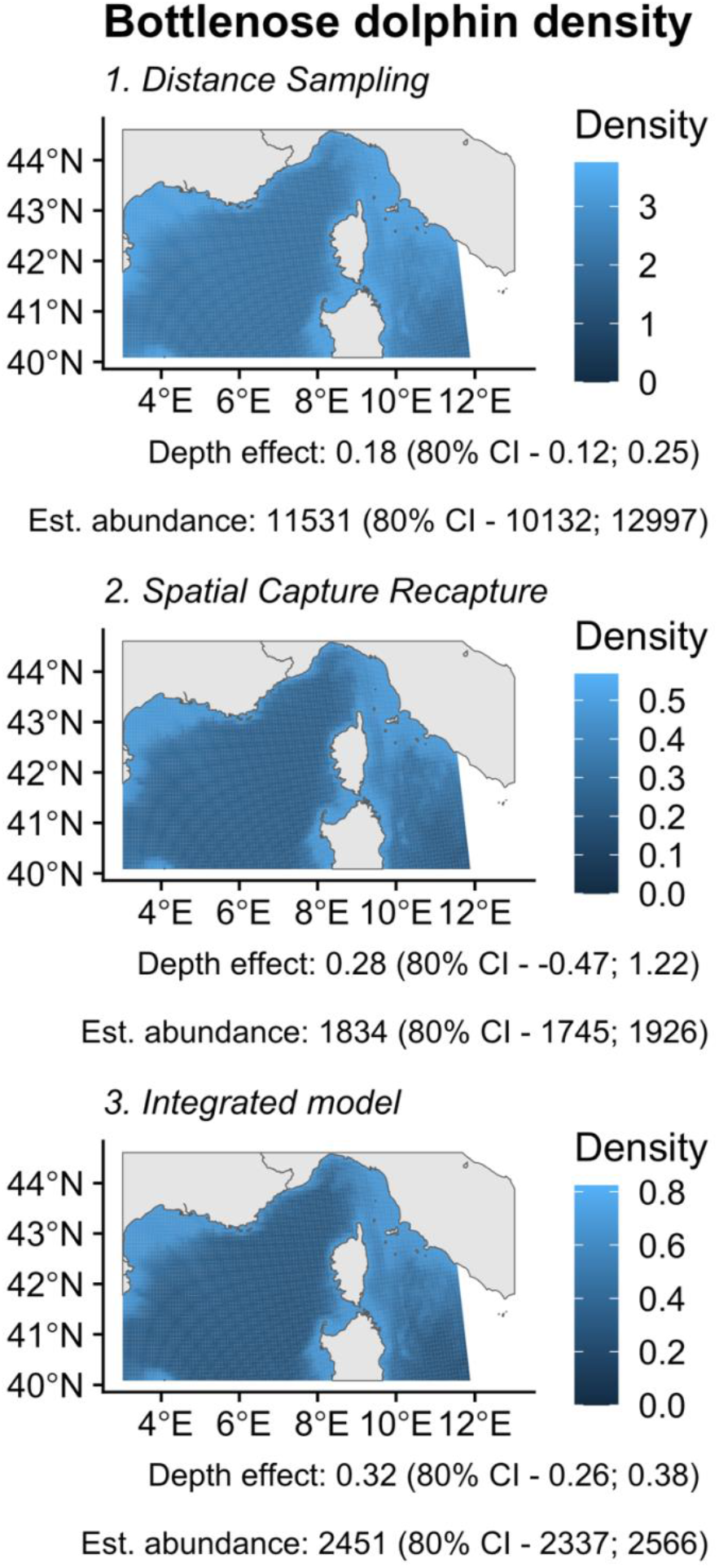
**Density of bottlenose dolphins** (*Tursiops truncatus*) estimated from 1. Distance Sampling (DS), 2. Spatial Capture Recapture (SCR), 3. Integrated model. Lighter colour indicates more individuals per area unit. All models predicted higher density in coastal seas, while depth effect is no significant for SCR model. Note that density scales are different between maps, indicating a higher overall population size for DS model than for Integrated model and SCR model.

### Conservation implications for monitoring bottlenose dolphins in the French Mediterranean Sea and beyond

When the conservation goal is to assess abundance in an area at a specific time, line transect surveys may be a cost-effective choice. However, if one’s goal is to estimate the number of animals in an area over a longer period, CR methods could be more appropriate but have cost implications that may exceed those of conducting a line transect survey (Crum, Neyman, & Gowan, 2021; Hammond et al., 2021). Despite differences in ecological inference, DS and CR are complementary methods depending on the conservation motivations and funding. To date, assessing bottlenose dolphin population of French Mediterranean Sea for the EU reporting only focuses on the DS data (Laran et al., 2017). Aerial surveys provide crucial information on marine megafauna taxa, and on human pressures to fill several criteria of the Marine Strategy Framework Directive (Laran et al., 2017; Pettex et al., 2017; Lambert et al., 2020). However, funding constraints make the aerial monitoring hardly applicable at a high frequency, and it is planned to be implemented every 6 years. In parallel, the French office for biodiversity develops and supports local monitoring programs in the French MPA network to perform photo-id data continuously, such detailed datasets represent an important asset to inform abundance of marine mammals populations (Evans & Hammond, 2004). Ecological indicators required by the Marine Strategy Framework Directive for bottlenose dolphins would benefit from integrating aerial line-transect with more data when available (Lauret et al. 2021). In addition, the French Research Institute for Exploitation of the Sea (i.e. IFREMER) collected yearly bottlenose dolphins’ data during line transects surveys for pelagic fisheries (Baudrier et al., 2018). Ultimately, several monitoring programs will be available about bottlenose dolphins in the Mediterranean context and integrated spatial models makes possible to include existing datasets that have been discarded so far to inform public policies (Cheney et al., 2013; Isaac et al., 2019).

We acknowledge that our model has limitations due to several ecological features lacking, e.g. spatial autocorrelation, effect of other environmental covariates, accounting for non-linear covariate effect, and group behaviour of bottlenose dolphins that may generate non-independent individual detection probabilities. One might also consider extending the activity center process to include a movement model of individuals (Gowan, Crum, & Roberts, 2021). Moreover, ecological closure assumptions we assumed are likely to be violated but we assumed the bias introduced bias would be minimal. However, we emphasize that integrated spatial models are highly relevant considering the future monitoring planning by the French biodiversity agency that will perpetuate the coexistence of photo-identification with aerial line-transect. Analysing the collected data in an integrated framework will lead to a more comprehensive understanding of how the monitoring programs can work together and what exactly it is that they achieve in unison. It is our hope that the ability of integrating different datasets contribute to the ongoing monitoring efforts developed in the Mediterranean context and fit in the scope of what managers expect form statistical developments to inform environmental policies (Lauret, 2021). Line-transect and capture-recapture surveys are widely used monitoring methods to assess population dynamics of marine mammals (Hammond et al., 2021). Our work provides a promising modelling baseline to deal with the bottlenose dolphin evaluation but also open perspectives for other conservation challenges about marine species that are subject to similar monitoring situations in the French Mediterranean context (e.g. fin whale, seabirds) and elsewhere.

Last, adding complementary long-term datasets to the aerial-surveys would make possible to access the demographic parameters (*e*.*g*. recruitments, survival (Chandler et al. 2018)), which would represent a major opportunity for the knowledge about French Mediterranean bottlenose dolphin populations and to produce reliable conservation status. The use of integrated spatial models for the French Mediterranean bottlenose dolphin population also enable to extend the modelling approach exploring seasonality in density, and to measure immigration and dispersal between bottlenose dolphins populations (Zipkin & Saunders, 2018). Finally, precising the assessment of bottlenose dolphin conservation status could ultimately lead to mitigation programs in the context of the Marine Strategy Framework Directive, e.g. marine protected areas implementation such as the Bottlenose dolphins Natura 2000 area in the French Gulf of Lion.

### Spatial integrated models as a promising tool for conservation

When establishing species conservation status for large-scale environmental policies, discarding some datasets from the analysis can reduce the reliability of the ecological estimation (Bischof, Brøseth, & Gimenez, 2016). Using multiple datasets into integrated spatial models help to overcome some limitations present when using separated information sources (e.g. limited spatial or temporal survey coverage, Zipkin & Saunders 2018; Isaac et al. 2019). However, caution should be taken as integrating data requires additional modelling assumptions, e.g. assuming population closure over longer time period in our case (Dupont et al., 2019; Farr et al., 2020; Fletcher et al., 2019; Simmonds et al., 2020). Integrated spatial models are flexible tools that can include more than 2 datasets (Zipkin & Saunders, 2018), and various type of data that enlarge the scope of usable information (presence-absence (Santika et al. 2017), count data (Chandler et al., 2018), citizen science data (Sun, Royle, & Fuller, 2019)). Recent and current developments of SCR models widen perspectives to extend integrated spatial models to account for unidentified individuals, or to better describe animal movement (Milleret et al., 2019; Jiménez et al., 2020; Turek et al., 2020). Over the last decades, the spatial scope of conservation efforts has greatly increased, and the analytical methods have had to adapt accordingly (Zipkin & Saunders, 2018). Integrated spatial models are a promising tool that can be used in multiple situations where several data sources coexist, especially for large scale conservation policies.

## ACKNOWLEDGEMENTS

The French Ministry in charge of the environment (Ministère de la Transition Energetique et Solidaire) and the French Office for Biodiversity (OFB) funded the project SAMM that performed the aerial line-transects. The PELAGIS observatory, with the help of the OFB, designed, coordinated and conducted the survey: Emeline Pettex, Ghislain Doremus and Olivier Van Canneyt. We thank all the observers: Léa David (cruise leader), Eric Stéphan (cruise leader), Thomas Barreau, Ariane Blanchard, Vincent Bretille, Alexis Chevalier, Cécile Dars, Olivier Dian, Nathalie Di-Méglio, Emilie Durand, Marc Duvilla, Emmanuelle Levesque, Alessio Maglio, Marie Pellé, Morgane Perri, and Sandrine Serre. We are indebted to all aircraft crew members of Pixair Survey and the logistic partnership SINAY. We are grateful to all financial partners of the GDEGeM project that performed the photo-identification monitoring. We warmly thank technical and scientific participants of GDEGeM.

## Notes

https://github.com/valentinlauret/SpatialIntegratedModelTursiops

### Competing Interest Statement

The authors have declared no competing interest.

### Summary of Updates

Upload minor modifications according to review in a journal

