## Appendix 1 for "Integrated spatial models foster complementarity between monitoring programs in producing large-scale bottlenose dolphin indicators" (from another version of this preprint: version 2, 2022-04-15)

### Appendix 1: Data exploration: Spatial Integrated Model

Valentin Lauret, Hélène Labach, Daniel Turek, Sophie Laran, Olivier Gimenez

---

In this document, we provide data exploration and results of our analyze of bottlenose dolphins *(Tursiops truncatus)* in the northwestern Mediterranean Sea. This is the appendix 1 of our manuscript entitled *Integrated spatial models foster complementarity between monitoring programs in producing large-scale bottlenose dolphin indicators*, preprint available https://doi.org/10.1101/2021.02.01.429097.

To study bottlenose dolphins, we combined two existing datasets in the French waters :

- aerial line-transects collecting bottlenose dolphins data following a distance sampling (DS) protocol.
- at-sea photo identification collecting indivdiual data about dolphins population.

We built a DS model to analyze aerial data, a spatial capture-recapture (SCR) model to analyze at-sea data, and a spatial integrated model (SIM) to analyze jointly both datasets and to estimate abundance and density.

Hereafter, we provide data exploration and displayed some of the results.

#### The Data

The objective of this section is to provide supplementary information about bottlenose dolphins data.

#### About bottlenose dolphins

Bottlenose dolphins *(Tursiops truncatus)* in the North-Western Mediterranean Sea. In the marine world, many species of conservation interest are elusive, and ecological data can be costly to obtain. In particular, the high seas are difficult to access and ecological monitoring is often performed through aerial surveys. However, coastal seas allows performing detailed at-sea monitoring. Besides, many species such as marine megafauna are mobile and occur in both coasts and high seas. Combining monitoring programs that are carried out in each realm (i.e. coasts and high seas) has the potential to provide relevant information about these species.

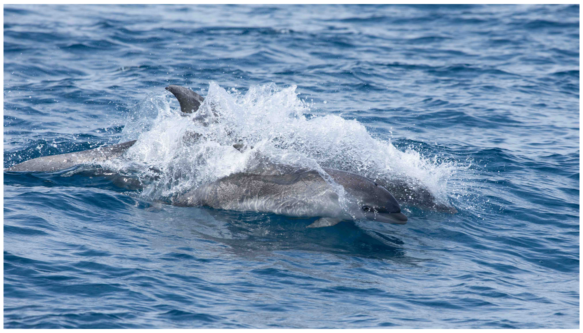

Figure 1: Bottlenose dolphins in the French Mediterranean Sea

### Monitoring programs

We focused on the North-Western Mediterranean, an area of 255,000 km2, which includes the Gulf of Lion and the Ligurian sea, the French coast of Provence, Corsica, and the Northern part of Sardinia (Figure 2). Our study area includes the Pelagos Sanctuary, which is a transboundary marine protected area for Mediterranean marine mammals covering an area of 90,000 km2 between Italy, France and Monaco.
The North-Western Mediterranean Sea is a critical habitat for many cetaceans species. Due to its coastal behaviour, bottlenose dolphins suffer from several anthropogenic pressures (e.g. collisions, fisheries bycatch, pollution, or acoustic perturbations), which raise concerns about their coexistence with human activities. The Mediterranean population of Bottlenose dolphins is considered "vulnerable" by the IUCN Red List and is one of the two cetacean species listed on Annex 2 of the European Habitats Directive (92/43/EEC). The protected status of this species within the French seas led to the development of specific programs to monitor Mediterranean bottlenose dolphins within the implementation of the European Marine Strategy Framework Directive (2008/56/EC; MSFD).

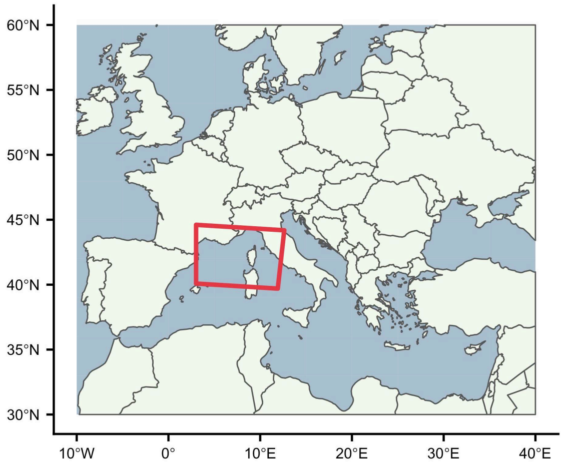

Figure 2: Location of study area in the Mediterranean basin.

#### At-sea monitoring

We used data from the first large-scale study of Bottlenose dolphins in the French Mediterranean Sea. Four NGOs and one marine reserve performed at-sea surveys over 21,464 km of the French continental shelf including the Gulf of Lion, the French Riviera, and Corsica (Figure 2). Observers performed monitoring aboard small sailing and motor boats to locate and photo-identify bottlenose dolphins all year long between 2013 and 2015 (observers collected group size, behaviour, and took pictures of the dorsal fin of each individual in the group when possible). Such surveys with small vessels are expected to recorded detailed information while being limited to coastal area. Between summer 2013 and summer 2015, at-sea surveys detected 129 bottlenose dolphin groups located in 89 different grid-cells. The sampling effort of at-sea surveys was heterogeneous in space (i.e. between 1 and more than 500 km prospected per sampled grid-cell), and time (i.e. higher prospection in spring and summer than in autumn and winter). At-sea surveys did not include repeated visits. Some sites have been visited once, and others have been visited 50 times.

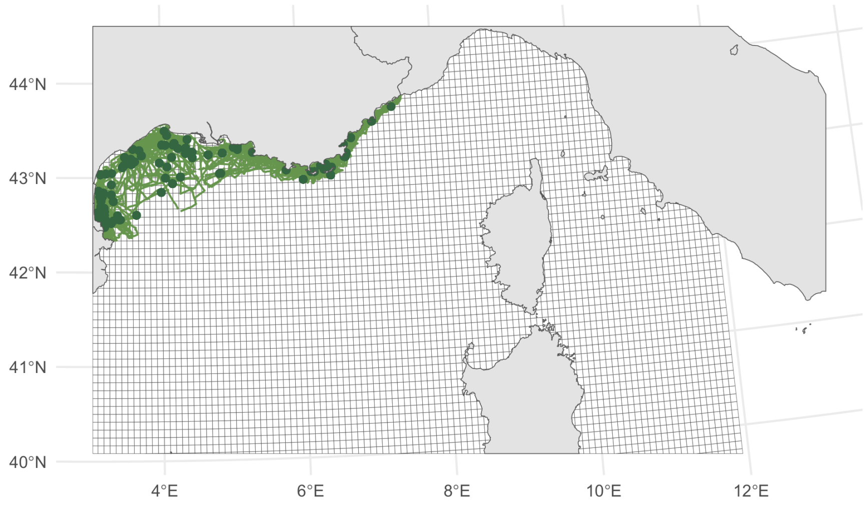

Figure 3: Boat surveys (GDEGeM program) prospected 21,646 km of of the French continental shelf. Darkgreen dots are detection locations.

#### Aerial pelagic and coastal monitoring program

Data were collected during aerial surveys targeting the main taxa of marine megafauna within the French Exclusive Economic Zone (EEZ) including the Pelagos Sanctuary. The survey covered 24,624 km of line-transect performed by scientific institutional partners of the French Biodiversity Office and sampled 1336 grid-cells (i.e. 30.67% of the total number of grid-cells, Fig. 1). Two trained observers collected cetacean data following a distance sampling protocol (i.e. recording species identification, group size, declination angle). Aerial surveys were conditional on a good weather forecast and were performed using high-wind aircrafts with bubble windows. Sampling effort for aerial surveys was homogeneous over the studied area with three or four replicates per line-transect between November 2011 and August 2012. While aircrafts surveyed large area quickly, the limited temporal coverage may reduce overall number of detections. In the future, this survey will be conducted every six years to inform the MSFD (for more details).

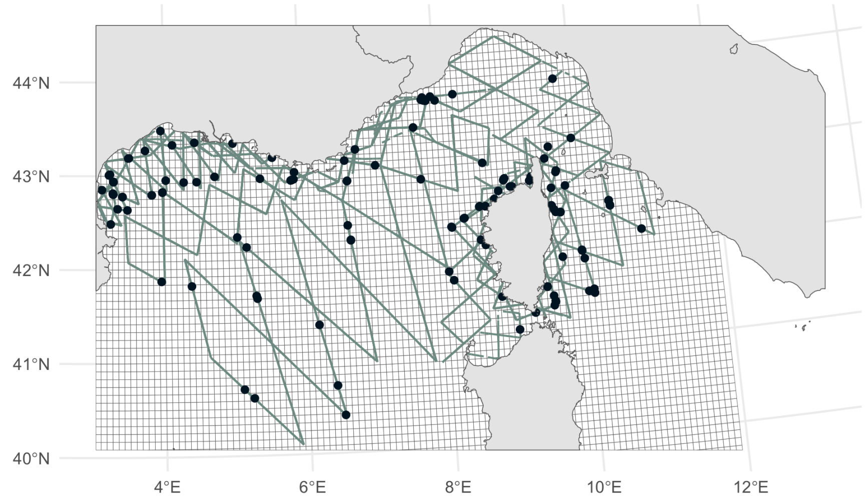

Figure 4: Aerial surveys (SAMM program) prospected 24,624 km of both sea shelf and high seas, darkblue dots are detection locations.

The two monitoring programs collected either photo-identification or distance sampling data. For both programs, we used the locations of bottlenose dolphins encounters and the survey tracks. We used depth as an environmental covariate to estimate density of bottlenose dolphins.

### Results: comparison between models

#### Density maps

Density maps are built projecting $\lambda$ in every site of the study area. This is the Fig. 2 of the manuscript. We observed that all models displayed a positive effect of shallow waters on dolphin density.

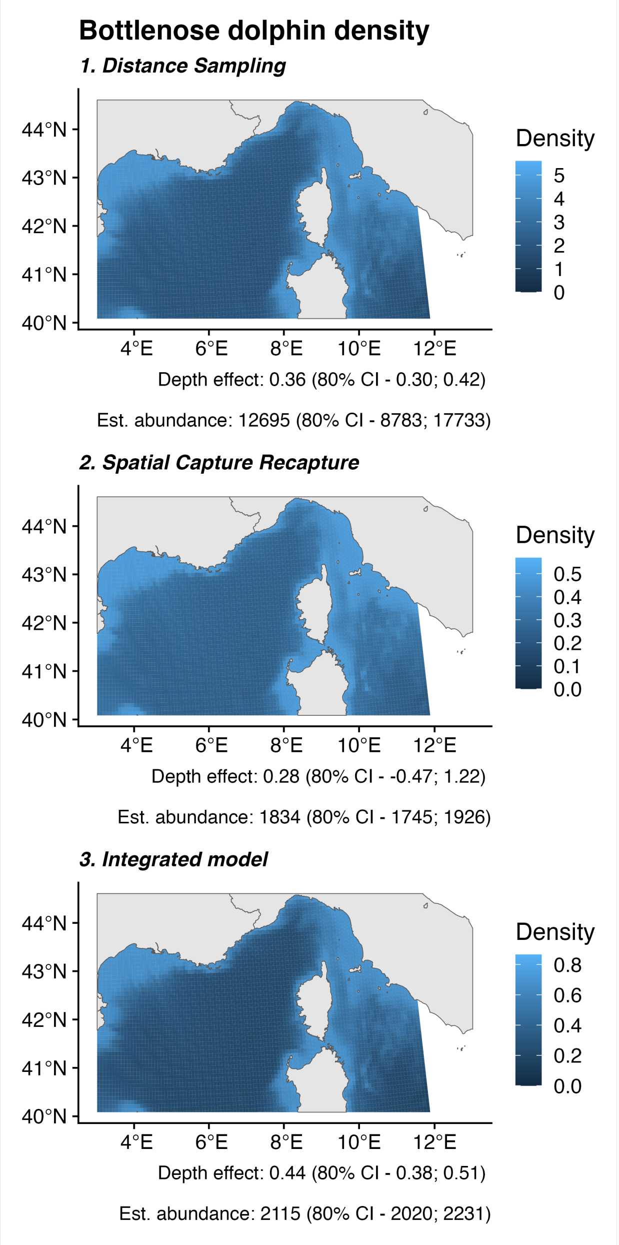

Figure 5: Density of bottlenose dolphins estimated by DS model, SCR model, and SIM

#### Differences of density

We displayed a couple of plots that illustrate the pixel-by-pixel relative density difference between IM-SCR and IM-DS (just simple raster algebra). DS model estimates higher density than IM, especially in the pelagic seas whereas IM predicted higher densities than SCR model thrgouhout the area.

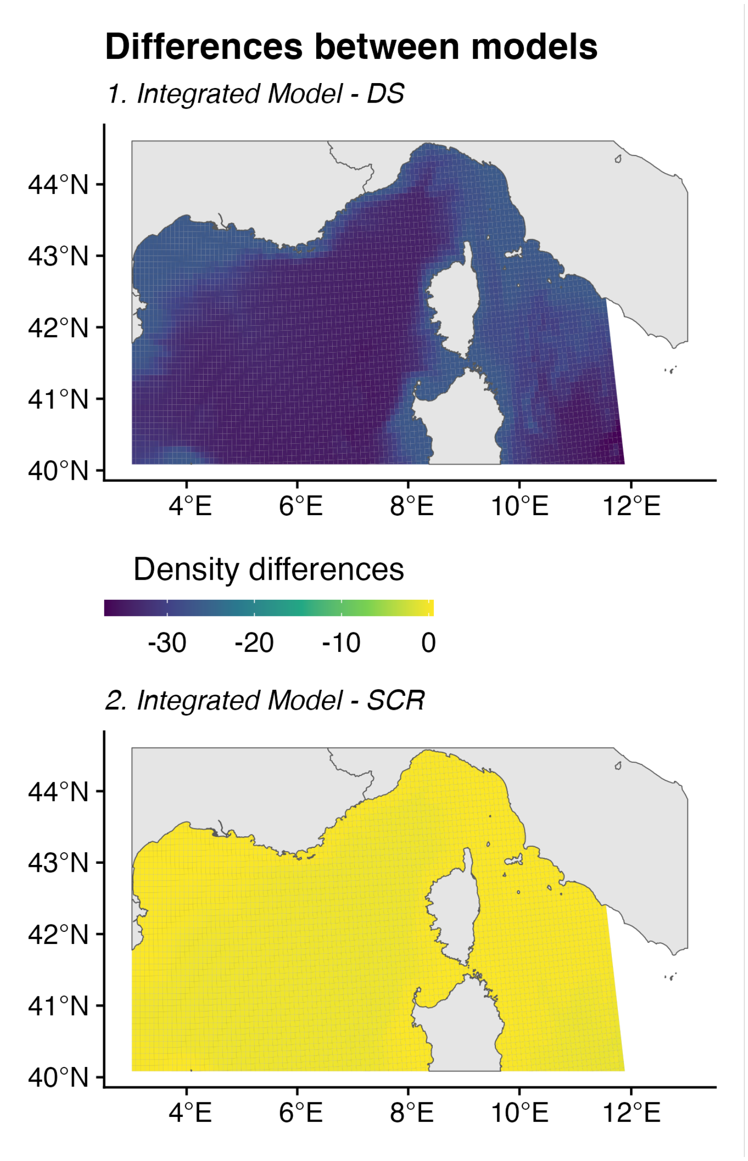

Figure 6: Differences between model estimates. 1. Density estimated by DS substracted to density estimated by the Integrated model, 2. Density estimated by SCR substrated to density estimated by the Integrated model. Negative differences indicate that DS (or SCR) model predicted more dolphins than the integrated model. Positive differences indicate that the integrated model predicted more dolphins than DS (or SCR).

##### Maps of activity centers

Both models are predicted a coastal repartition of activity centers although we observed that IM predicted more pelagic activity centers than SCR model.

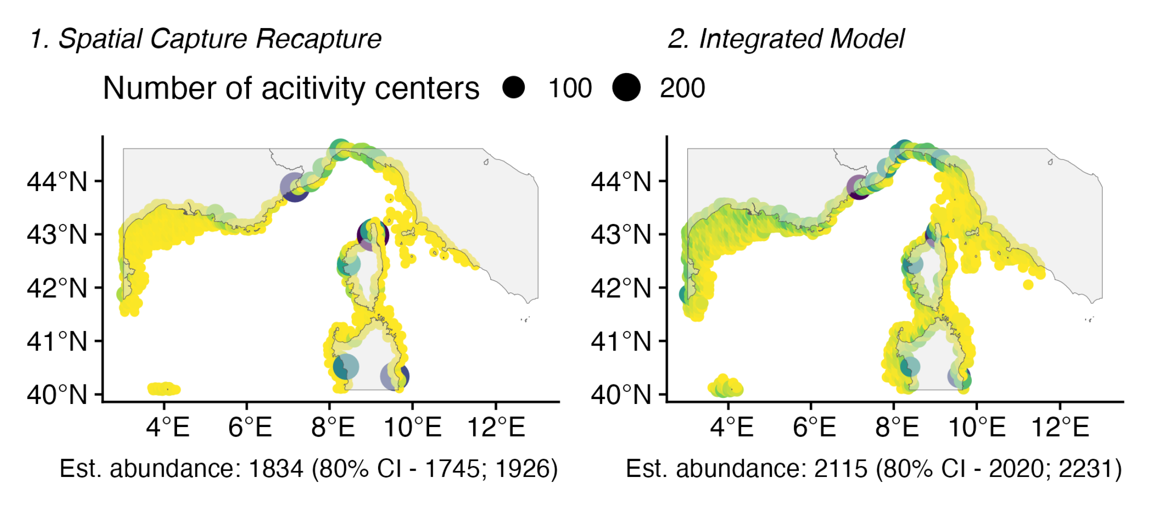

Figure 7: Map of bottlenose dolphins activity centers estimated by 1. The Spatial capture recpapture model, 2. The integrated model. We displayed bottlenose dolphin abundance estimated by each model. Fill color of circles reflected the number of dolphins activity centers assign to each location.
